# The Cation/H^+^ exchanger OsCAX2 is involved in cadmium tolerance and accumulation through vacuolar sequestration in rice

**DOI:** 10.1101/2022.11.22.517486

**Authors:** Wenli Zou, Junhui Zhan, Lijun Meng, Yuetong Chen, Dandan Chen, Mingpei Zhang, Haohua He, Jingguang Chen, Guoyou Ye

## Abstract

Excessive cadmium (Cd) in rice grains is a serious food safety problem. The development of Cd-safe varieties requires the identification of germplasms and genes with major effect on Cd accumulation but without negative effects on other important traits. Here, we reported that OsCAX2, a member of the rice Cation/H^+^ exchanger (CAX) family, is an important Cd transporter. *OsCAX2* encodes a tonoplast-localized protein and is strongly upregulated by Cd, mainly expresses in root exodermis, parenchyma in cortex, endodermis and stele cells. Depletion of *OsCAX2* resulted in enhanced Cd sensitivity and root-to-shoot translocation in rice, while overexpression of *OsCAX2* significantly increased Cd tolerance and reduced Cd transport by promoting root Cd influx and vacuolar storage, which ultimately reduced Cd transport via xylem. *OsCAX2* also had significant effects on tissues/organs distribution of Cd but had no effects on grain yield and agronomic traits. Importantly, the *OsCAX2* overexpressing lines had more than 70% lower grain Cd accumulation, increased iron (Fe), zinc (Zn) and manganese (Mn) and reduced copper (Cu) accumulation. Therefore, *OsCAX2* is an ideal gene for developing Cd-safe rice varieties via transgenic approach.

## 1 INTRODUCTION

Cadmium (Cd) is one of the most toxic heavy metal elements in the environment. Cd stress has an inhibitory effect on seed germination, photosynthesis rate, enzyme activity, and seriously affects the growth and development in plants (Nagajyoti et al., 2010; Sharma and Agrawal, 2005; Zhu et al., 2021). Excessive Cd in rice grain is a major food safety issue, since Cd has serious harmful effects on human health and rice is the staple food of more than half of the world population (Chen et al., 2019). Thus, it is imperative to reduce Cd accumulation in brown rice

Plants as an adaptive strategy have developed efficient physiological and biochemical mechanisms to counter Cd stress, including reducing Cd bioavailability, decreasing Cd influx, promoting Cd efflux, sequestrating Cd into vacuole, chelating Cd in cytosol, and detoxification of Cd-induced reactive oxygen species (ROS) (Baliardini et al., 2015; DalCorso et al., 2010; Ebbs et al., 2009; Lin and Aarts, 2012; Luo and Zhang, 2021; Ye et al., 2012). Membrane-localized transporters are the most important in Cd homeostasis by involving in uptake, transport and accumulation in rice. Cd is first transported into plants through carrier proteins in roots, most of which are localized on plasma membrane. The plasma membrane located OsNRAMP5 is a key transporter responsible for Cd uptake in roots (Chang et al., 2020; Ishimaru et al., 2012). Knockout of *OsNRAMP5* results in the loss of Cd uptake capacity in rice roots (Chang et al., 2020), and significantly reduces Cd content in flag leaves and grains (Yang et al., 2019). However, significant negative effects on grain yield and other agronomic traits have been reported for some of the *osnramp5* mutants, which is likely caused by the significant decrease in Mn accumulation in the mutants (Yang et al., 2019). OsNRAMP1, highly homologous to OsNRAMP5, is located on plasma membrane and promotes Cd uptake in roots (Takahashi et al., 2011). Overexpression of *OsNRAMP1* enhances Cd accumulation in roots and shoots (Takahashi et al., 2011). The plasma membrane located OsCd1 positively contributes to Cd uptake and transport in rice (Yan et al., 2019). The root-to-shoot translocation and distribution to different tissues are key steps for Cd distribution to the grains. The plasma membrane located OsHMA2 and OsCCX2 play important role in the Cd translocation from roots to shoots with knockout mutations reducing grain Cd content and yield or seed weight (Hao et al., 2018; Satoh-Nagasawa et al., 2012; Takahashi et al., 2012; Yamaji et al., 2013). The tonoplast-localized OsHMA3 and OsABCC9 sequestrate Cd into vacuoles and restrict Cd translocation from roots to shoots (Cai et al., 2019; Yang et al., 2021). The natural variation of *OsHMA3* greatly contributes to the different grain Cd contents of rice varieties and subspecies with functionally defective mutations resulting in increased root-to-shoot transport and grain Cd accumulation (Ueno et al., 2009; Yan et al., 2016). So far, no adverse effect on grain yield and other agronomic traits has been reported for *OsHMA3*, probably because it has a relatively high Cd specificity. The existence of natural variants also makes *OsHMA3* applicable in conventional breeding, although desirable alleles (reducing grain Cd content) with larger effects than the common allele in japonica varieties are yet to be identified. The tonoplast-localized OsNRAMP2 mediates Fe and Cd efflux from vacuole to cytosol (Chang et al., 2022; Li et al., 2021). The endoplasmic reticulum (ER)-located OsLCT2 limits the xylem loading of Cd and restricts the root-to shoot translocation by sequestrating Cd into ER (Tang et al., 2021). The plasma membrane located OsLCT1 is involved in transporting Cd to the grains likely through the xylem to phloem transfer process and its mutation significantly reduces grain Cd accumulation (Uraguchi et al., 2011). *OsHAM2* also contributes to Cd distribution to grains via nodes (Takahashi et al., 2012; Yamaji et al., 2013).

The Cation/H^+^ exchanger (CAX) transporters have been shown to play important roles in Cd tolerance and accumulation in *Arabidopsis thaliana*, and the Cd hyper-accumulators *Arabidopsis halleri* and *Sedum alfredii* (Baliardini et al., 2015; Koren’kov et al., 2007; Korenkov et al., 2007; Zou et al., 2021). Overexpression of *SaCAX2h, AtCAX2* and *AtCAX4* enhances Cd tolerance and improves root Cd sequestration in transgenic tobacco (Koren’kov et al., 2007; Korenkov et al., 2007; Korenkov et al., 2009; Zhang et al., 2015). Loss-of-function of *AhCAX1* results in a high sensitivity to Cd but only at low concentration of Ca (Baliardini et al., 2015). The *ahcax1* and *atcax3* mutants have higher Cd sensitivity and stronger ROS accumulation in *Arabidopsis thaliana* (Ahmadi et al., 2018; Baliardini et al., 2015; Modareszadeh et al., 2021). We recently showed that a few members of the rice CAX family have Cd transport activity in yeast (Zou et al., 2021). However, whether and how the rice CAX transporters involve in Cd tolerance and transport in rice has not been reported yet.

In this study, we functionally characterized the role of *OsCAX2* in Cd tolerance and accumulation in rice. Our results suggested that *OsCAX2* is a Cd transporter and significantly contributes to tolerance and accumulation of Cd in rice mainly through involving in root uptake, vacuolar sequestration, xylem loading and tissue distribution of Cd and that overexpression of *OsCAX2* can be used to produce Cd-safe rice lines without adverse effects on yield, agronomical traits and contents of essential metal elements.

## 2 MATERIALS AND METHODS

### 2.1 Plasmid construction and plant transformation

To generate the *CRISPR-oscax2* knockout (KO) mutant and overexpression of *OsCAX2* (OE) lines, the sgRNA-Cas9 and *pCAMBIA1300-OsCAX2* plant expression vectors were constructed. The oligos corresponding to the guide RNA sequences of *OsCAX2* were synthesized and cloned into the CRISPR/Cas9 genome editing vector as previously described (Meng et al., 2017). The coding DNA sequences of *OsCAX2* (1317 bp) were cloned into the pEASY-Blunt vector (TransGen Biotech, Beijing), were then inserted into pDONR221 via BP reactions to generate entry vectors, followed by LR reactions (Invitrogen, USA) with *pCAMBIA1300-pUbi-GFP* or *pCAMBIA1300-pUbi*. Nipponbare (*Oryza sativa* L. ssp. japonica) was used as the hosts in agrobacterium-mediated transformation for producing the KO and OE lines. To identify the mutations of the KO lines, mature seeds were collected from the T_0_ plants and geminated for genotyping. Total DNA isolated from the transgenic plants was used to amplify the genomic region surrounding the CRISPR target sites using specific primers, and the PCR products were directly sequenced using *OsCAX2* specific primers by Sanger method. All plasmid constructions were confirmed by sequencing (Sangon Biotech, Shanghai). T_2_–T_3_ generations of homozygous transgenic lines were used in the experiments described below. All primers used for vector construction are listed in Supplemental Table 1.

### 2.2 Plant materials and growth conditions

The rice variety Nipponbare used as wild type (WT) and three KO and three OE lines were used in this study. The basic hydroponic culture procedure was as follow. Seeds were surface-sterilized and germinated with deionized water for 48 h in the dark at 30 °C. The well germinated seeds were transferred into 96-well germinating plates without a base in a 3.5 L plastic pot filled with deionized water for 7 days. The seedlings were then cultured in nutrient solution prepared according to the composition of the International Rice Research Institute (IRRI) solution (Hou et al., 2021). HCl/NaOH (1 M) was used to adjust the pH, and 0.39 g L^-1^ C_6_H_13_NO_4_S (MES) was added to stabilize the pH at 5.5. Culture conduction in the growth chamber was: 16 h/8 h day/night, temperature cycle of 30 °C/25 °C, 800 μmol m^-2^ s^-1^ light intensity and 60–65% relative humidity. The solution was changed every 48 h. The seedlings with uniform growth were used in experiments described below.

To investigate the effects of *OsCAX2* on Cd tolerance and accumulation at seedling stage, two-week old seedlings were exposed to the IRRI solution containing 0, 0.1, 1, 2 or 5 μM CdCl_2_ for 7 days. A randomized complete block design with three 12-plant replicates was adopted for all the treatments. After treatment, plants were sampled and roots and shoots were harvested separately for growth assay and Cd determination.

A pot experiment was carried out in the greenhouse of the Agricultural Genomics Institute in Shenzhen, Chinese Academy of Agricultural Sciences (AGIS-CAAS, Shenzhen, China) from July to November, 2020. The soil was collected from an uncontaminated paddy field (total Cd 0.1 mg kg^-1^, pH at 6.0~6.2) and amended with 1.5□mg□kg^-1^ Cd. Each pot was filled with 15 kg soil. Three-week-old seedlings of the WT, KO and OE lines pre-cultured hydroponically were transplanted into pots with 3 plants per pot. A randomized complete block design with six one-pot replicates was used to layout the experiment. All pots were irrigated with distilled water every day to maintain a water level of 3 cm above the soil surface. The tissue/organ samples were taken at maturity for measuring contents of Cd and other metal elements.

A field experiment was conducted in a Cd-polluted paddy field at AGIS-CAAS (Shenzhen, China) from June to October, 2021. The soil Cd concentration is 1.8 mg kg^-1^ and pH at 5.5. A randomized complete block design with two replicates and a plot size of five 5-plants rows was used to layout the experiment. The local cultivation practices were followed to manage the experiment properly. The tiller development and grain yield were observed at maturity stage. For the measurement of metal concentrations, the tissue/organ samples were taken at maturity. For measuring the agronomic traits and grain yield, the well-established practices were followed.

### 2.3 Xylem sap collection assay

To determine the Cd accumulation in the shoot xylem sap, three-week-old hydroponically grown seedlings were exposed to IRRI solution containing 5 μM CdCl_2_ for 3, 6, or 9 days. The shoots were cut off at 2 cm above the roots using a razor blade and the xylem sap was collected for 2 h using a micropipette. The first drop was discarded to avoid contamination from damaged cells. Xylem sap collected from 12 plants was pooled into one replicate, and three replicates were used for each line. After centrifugation at 12,000 g for 10 min the samples were used to measure the content of mineral elements as described below.

### 2.4 Cd uptake kinetics experiment

To assess the uptake of Cd, a short-term uptake experiment was conducted. Three-week-old hydroponically grown seedlings were pretreated in the uptake solution (0.5 mM CaCl_2_, 2 mM MES, pH at 5.6) at room temperature for 12 h (Yan et al., 2016) and then transferred to the uptake solution containing different Cd concentrations (0, 0.2, 0.5, 1, 2 or 5 μM CdCl_2_) at 4 °C and 25 °C for 20 min under continuous aeration. At the end of the experiment, the roots were washed in ice-cold uptake solution without Cd for 5 min and rinsed with deionized water three times. The root samples were collected by separating the roots from the shoots with a sharp blade and dried at 70 °C for 3 days. Three biological replicates with at least 6 plants per replicate per line were used. The time-dependent Cd uptake kinetics experiment was conducted similarly with 5 μM CdCl_2_ treatment for 0, 1, 4, 12, 24, 48 or 72 h.

### 2.5 Transport activity assay in yeast

The heterologous yeast assay was utilized to examine the Cd and Ca transport ability of *OsCAX2*. The cDNA fragment containing an entire ORF of *AtNRAMP1, OsCAX2* and its two truncated versions (*OsCAX2-16AA* and *OsCAX2-40AA*) were amplified from the cDNA of rice variety Nipponbare and *Arabidopsis thaliana* ecotype Columbia-0 (Col-0). Using BamH I and EcoR I, the amplified gene sequences were ligated into the pYES2 vector (Invitrogen, USA), with correct direction, using a ClonExpress Ultra One Step Cloning Kit (Vazyme Biotech, Nanjing), generating the yeast expression vector. After sequencing confirmation, OsCAX2/pYES2 along with empty control, were individually transformed into the wild type *Saccharomyces cerevisiae* BY4741 (*MATa his3*Δ*1 leu2*Δ*0 met15*Δ*0 ura3*Δ*0*) and K667 (Δ*pmc1*Δ*vcx1*Δ*cnb1*, high Ca sensitivity of yeast mutant strain) yeast mutant strains according to the manufacturer’s protocol (Coolaber Technology, Beijing). Transformants were selected on synthetic dextrose medium without uracil (SD-Ura) containing 2% galactose, 0.67% yeast nitrogen base (Sigma-Aldrich), and 2% agar and verified by PCR with yeast plasmid extraction (Solarbio Technology, Beijing). Positive clones were cultured on SD-Ura liquid medium until the early logarithmic phase. The following experiments for yeast assay and analysis were all independently performed at least three times.

For evaluation of Cd tolerance inhibition, yeast cells of the BY4741 transformants were diluted to an OD_600_ of 1.0, 0.1, 0.01 and 0.001, step by step, with sterile water, and 6 μL of the cell suspension was spotted on SD-Ura plates containing 0, 40 or 80 μM CdCl_2_. The Ca tolerance assays were performed by culture the K667 transformants on solid yeast extract peptone dextrose (YPD) medium with 0, 50 or 150 mM CaCl_2_. The plates were incubated at 30 °C for 2 days before the growth phenotypes were evaluated.

To quantify the growth of the BY4741 transformants in liquid SD-Ura medium, yeast cells were cultured in SD-Ura liquid medium with 2% galactose at 30 °C with shaking at 200 rpm until the OD_600_ value reached to 0.8. Then 20 μL of cell suspensions were transferred to 20 mL fresh SD-Ura liquid medium with 10 μM CdCl_2_ and cultured overnight. The growth of the K667 transformants in liquid YPD medium was quantified in a similar way. Yeast cells were prepared and the OD_600_ was adjusted to 0.8 with sterile distilled water. Then 20 μL of cell suspensions was added to 20 mL fresh liquid YPD medium containing 0 or 100 mM CaCl_2_ and cultured overnight. The OD_600_ value was determined at the indicated time.

For determination of the Cd concentration of the BY4714 transformants and the Ca concentration of the K667 transformants, liquid culture was conducted as described above. The cultures of each set were harvested in pre-weighed microfuge tubes by centrifugation and washed with sterile water for three times. After aspirating the supernatant, pelleted cells were dried in the oven at 70 °C overnight.

To examine the subcellular localization of OsCAX2 in yeast, the OsCAX2-GFP fragment from the pCAMBIA1300-OsCAX2-GFP construct was cloned into the pYES2 vector to generate the pYES2-OsCAX2-GFP construct. The construct was introduced into the BY4741 yeast strain according to the method described above. Protoplasts were isolated from the OsCAX2-GFP yeast cells cultured overnight in darkness using a yeast protoplast preparation kit (Bestbio Biology, Shanghai). The fluorescence signal was observed by a TCS SP5 confocal laser scanning microscope (Leica Microsystems). All the images were further processed by using the Leica LAS AF Lite software (Leica LAS AF Lite, Germany).

### 2.6 Gene expression profile analysis

Total RNA was extracted from plant tissues using the TRIzol reagent (Vazyme Biotech, Nanjing). DNaseI-treated total RNAs were subjected to reverse transcription (RT) with the HiScript II Q Select RT SuperMix for qPCR (+gDNA wiper) kit (Vazyme Biotech, Nanjing). The transcript levels of selected genes were measured by RT-qPCR using the CFX96® Real □ Time PCR System (BioRad, http://www.bio□rad.com/) and the 2 × T5 Fast qPCR Mix (SYBRGreenI) kit (Vazyme Biotech, Nanjing) with the 2^-ΔΔCt^ method for data analysis. Three biological replicates were performed for RT-qPCR. The rice actin 1 gene (*Act1*) was used as an internal reference to normalize the gene expression. The primers for RT-qPCR are given in Supplemental Table S1.

### 2.7 Western blotting

The protein expression of OsCAX2 in rice leaves (1.5 g, three biological replicates) from the WT, KO and OE lines was confirmed by western blotting with an anti-GFP antibody. Protein extraction was conducted according to the method described previously (Shi et al., 2022). The actin was used as an internal control.

### 2.8 Cell and tissue specificity of *OsCAX2* expression

For histochemical analysis of *OsCAX2* expression, *pOsCAX2::GUS* transgenic line was generated as previously described (Zou et al., 2021). Samples of the *pOsCAX2::GUS* transgenic line subjected to designed treatments were collected and incubated at 37 °C for 24 h in the GUS staining solution (Coolaber Technology, Beijing). After staining, the green tissue materials were treated two or three times by 75% ethanol to remove chlorophyll and decolorize. GUS activity was detected by a stereoscopic fluorescence microscopy (TL5000, Leica Microsystems). For GFP observation of OsCAX2 tissue location, the promoter of *OsCAX2* was cloned into *pCAMBIA1300* and the *pOsCAX2::OsC*AX2-GFP vector was transformed into rice cultivar Nipponbare to generate transgenic plants. The roots of the transgenic and WT lines were collected and observed using a confocal laser scanning microscope at 500–535 nm after excitation at 488 nm.

Subcellular localization was investigated by transiently expressing OsCAX2-GFP fusion into tobacco (*Nicotiana benthamiana*) leaves and rice protoplasts isolated from the sheath of the WT seedlings (two-week-old) using the previously described method (Zhang et al., 2011). Double staining using AtTPK1 (red signal) as the tonoplast marker was used for further confirmation of the subcellular localization (Maitrejean et al., 2011). For co-localization experiments, sequential scanning was done for both of the channels and then merged together to show overlapping signals.

### 2.9 Visualization of Cd in plants

To avoid the interference of GFP signal, the OE lines without GFP-tagged, named OE# (OE1#, OE2# and OE3#) were used. The plant Cd fluorescent probe Leadmium™ Green AM (Invitrogen, USA) was used to monitor the localization of Cd in rice roots by following the manufacturer’s instructions. The properly treated samples were visualized and photographed using a fluorescence microscope with 488 nm excitation and 510–520 nm emission filters (Bahmani et al., 2019).

### 2.10 Protoplast Cd loading and fluorescence measurement

The protoplasts were isolated from the BY4741 yeast cells expressing *OsCAX2* or empty vector, as well as the WT, KO and OE# lines, then incubated in the presence of Leadmium™ Green AM for 60 min at 4 °C by the method described previously (Morel et al., 2009). After centrifugation, the pellet was re-suspended in a 0.85% NaCl solution. Then, the protoplasts were incubated in darkness for 5 min in the presence of 1 μM CdCl_2_, and the Cd localization and accumulation were detected using a confocal laser scanning microscope (Leica Microsystems). Fluorescence quantification was performed on 6 protoplasts for each sample.

### 2.11 Determination of Cd concentration in intact vacuoles

The determination of Cd concentration in intact vacuoles was performed according to the method described (Eroglu et al., 2016).

### 2.12 Measurement of metal concentration

All samples were oven dried at 70 °C for 3 days. The dried samples were crushed, wet-digested in concentrated HNO_3_ at 120 °C for 30 min, and further digested with HClO_4_ at 180 °C until the samples became transparent. The samples were then diluted with ultrapure water. The metal concentrations were determined using an inductively coupled plasma mass spectrometry (ICP-MS).

### 2.13 Statistical analysis

Statistical analysis was conducted using one-way ANOVA followed by planned multiple comparisons. *P* < 0.05 and *P* < 0.01 were regarded as significant and highly significant, respectively.

### 2.14 Accession numbers

The sequences of rice genes in this article can be downloaded from http://rice.plantbiology.msu.edu: *OsCAX2* (LOC_Os03g27960) and *OsActin* (AK100267).

## 3 RESULTS

### 3.1 Expression pattern and cellular localization of OsCAX2

To investigate the expression profile of *OsCAX2*, various tissues of the WT plants collected at seedling and flowering stages grown in a paddy field were analyzed via RT-qPCR. *OsCAX2* expressed highly in roots, moderately in nodes, leaf sheaths, leaves and flowering spikelet, and weakly in basal stems (Figure 1a). The response of *OsCAX2* expression to Cd treatment was investigated using hydroponically cultured seedlings. The expression of *OsCAX2* was elevated in roots and leaves in the presence of Cd, and increased with Cd dosage (0.1, 1, 2 or 5 μM), suggesting that *OsCAX2* was induced by Cd in both roots and leaves in a concentration-dependent manner (Figure 1b and c). Furthermore, in the time-course analysis of *OsCAX2* expression under 5 μM Cd treatment, the expression in roots was upregulated significantly at 0.5 h after treatment, and the maximum expression was observed at 4 h after treatment with about 7-fold increase compared to the control (Figure 1d), whereas the expression in leaves was significantly upregulated at 1 h after treatment, and the maximum expression was at 8 h after treatment with about 5-fold increase (Figure 1e). The expression of *OsCAX2* in roots increased with the distance to root tip in the presence of Cd (Figure 1f). These results indicated that *OsCAX2* is likely to involve in Cd stress response in rice.

**FIGURE 1.**
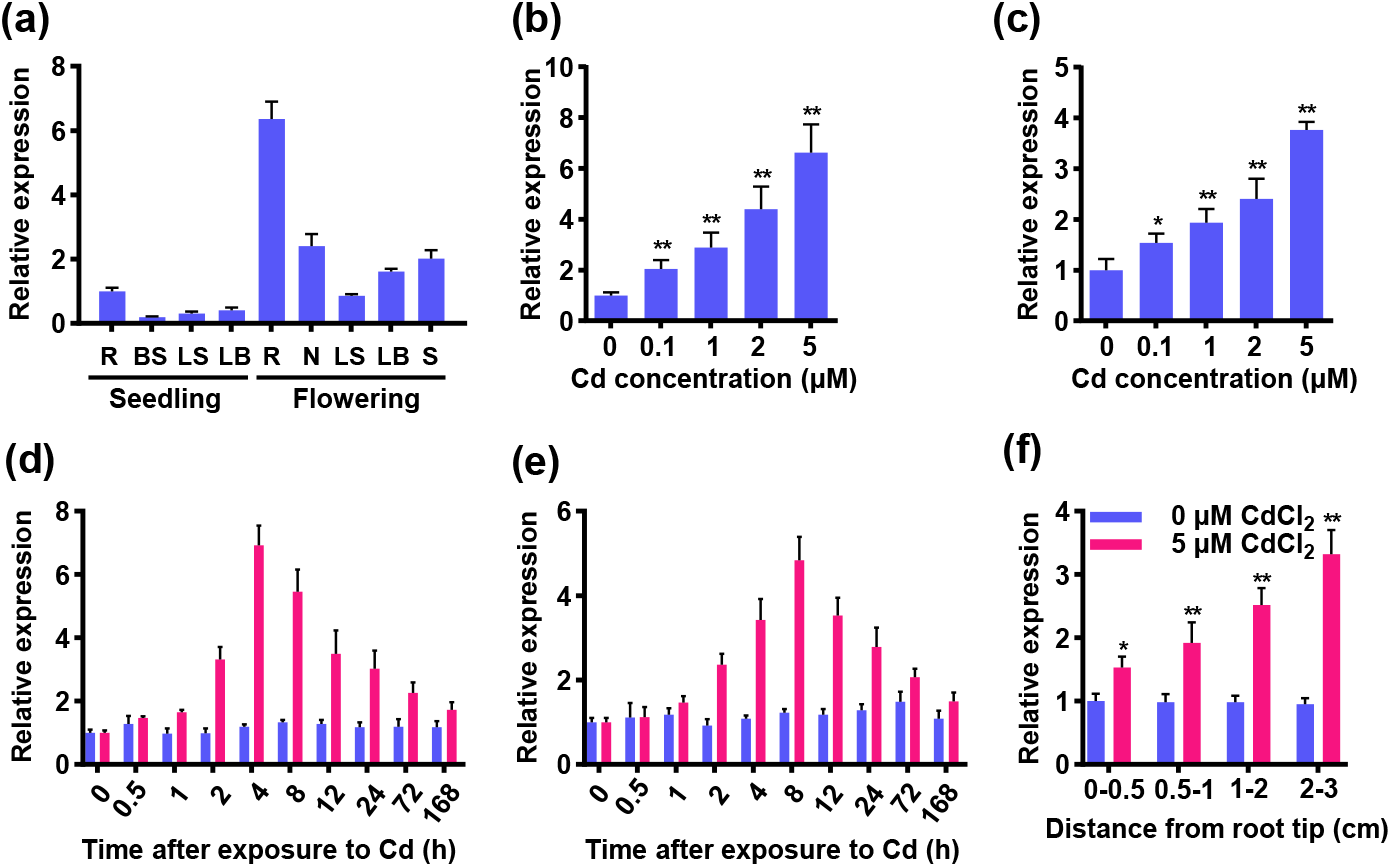
Expression analysis of *OsCAX2* in rice. (a) Expression levels of *OsCAX2* in tissues at different growth stage of Nipponbare (WT) grown in a paddy soil (total Cd 0.1 mg kg^-1^, pH at 6.0~6.2). The tissues are root (R), basal stem (BS), leaf sheath (LS), leaf blade (LB), node (N) and spikelet (S). (b-c) The expression levels of *OsCAX2* in the roots (b) and leaves (c). One-week-old hydroponically grown WT seedlings were exposed to treatments of different CdCl_2_ concentrations for 4 h. (d-e) Time-dependent expression of *OsCAX2* in the roots (d) and leaves (e). One-week-old seedlings were subjected to treatments (0 or 5 μM CdCl_2_) for at different durations. (f) Spatial expression pattern of *OsCAX2* in roots. Root segments 0-0.5, 0.5-1, 1-2 and 2-3 cm from the root tips of one-week-old seedlings exposed to 5 μM CdCl_2_ for 7 days. The rice actin 1 gene (*Act1*) was used as an internal control. Values are the mean ± SD of three independent replicates and statistical comparison was performed by ANOVA followed by planned comparison (\**P* < 0.05, \*\**P* < 0.01).

To investigate the tissue and cell specificity of *OsCAX2*, we generated transgenic lines expressing the GUS and GFP reporter driven by *OsCAX2* promoter. Histochemical staining of the GUS activities showed that *OsCAX2* expressed highly in roots, moderately in leaf sheaths, leaves and flowering spikelet (Figure 2a), consistent with the above RT-qPCR results (Figure 1a). The GFP signals of OsCAX2 were mainly detected in the root exodermis, parenchyma in cortex, endodermis and stele cells (Figure 2b).

**FIGURE 2.**
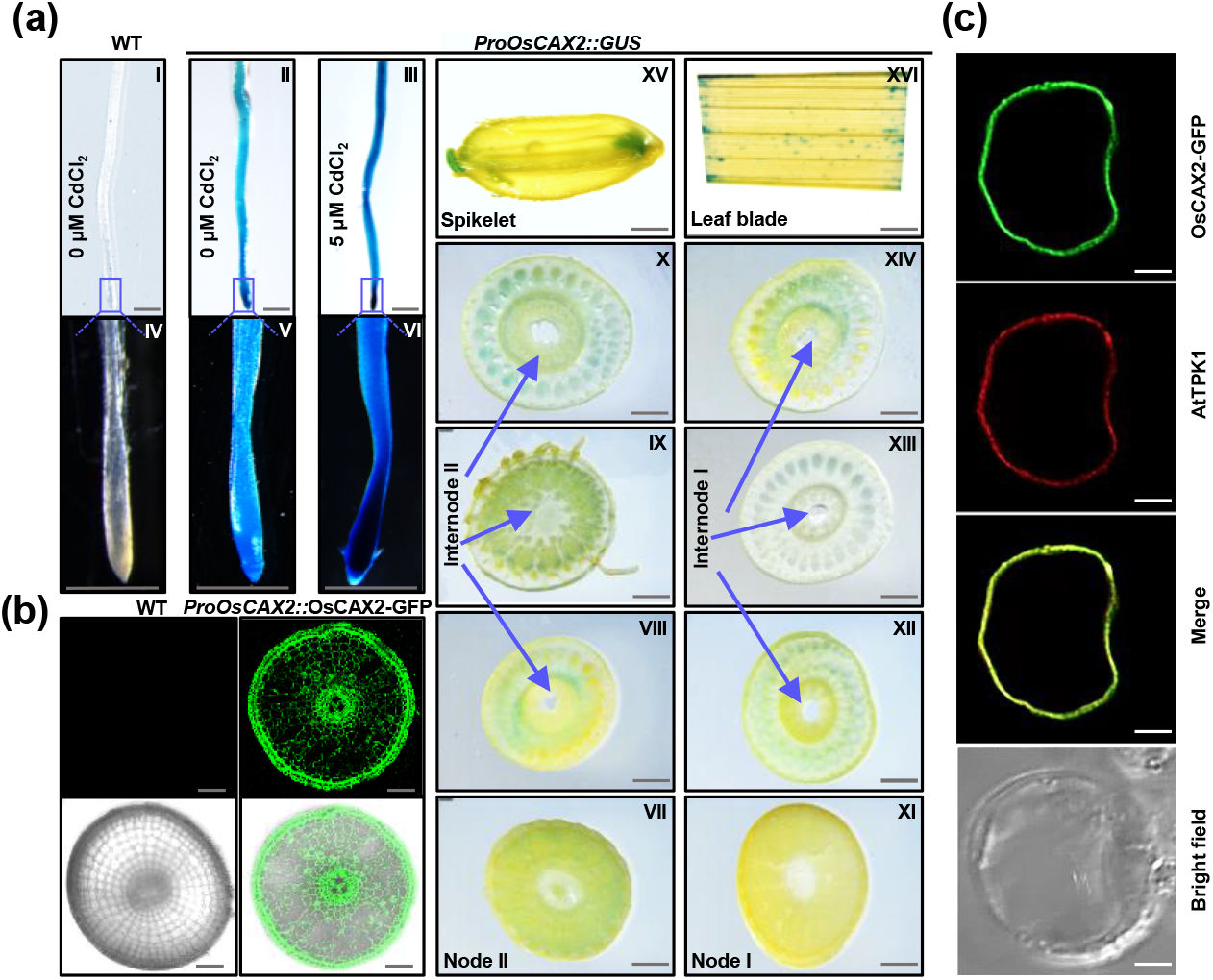
Tissue specificity and subcellular localization of OsCAX2 in rice. (a) Histochemical GUS assay of the transgenic plants with the GUS reporter driven by the *OsCAX2* promoter. I-III: GUS activity of the WT and *ProOsCAX2::GUS* in the roots exposed to solution with or without 5 μM CdCl_2_ for 7 days at seedling stage. IV-VI: magnified view of the root tips in I-III. VII-XVI: GUS activity of *ProOsCAX2::GUS* in tissues of field grown plants at flowering stage. The paddy soil has a total Cd content of 0.1 mg kg^-1^, and pH at 6.0~6.2. VII-X: Node II and internode II, XI-XIV: Node I and internode I, XV: Spikelet, XVI: Leaf blade. The blue arrows point to the position of internodes I-II. (Scale bar = 1 mm). (b) Tissue-specific localization of OsCAX2 in roots. WT and transgenic plants carrying the GFP driven by the *OsCAX2* native promoter (*pOsCAX2::OsCAX2-GFP*). WT, negative control. (Scale bar = 100 μm). (c) Subcellular localization of OsCAX2. The panels show the localization of OsCAX2-GFP and AtTPK1-RFP in rice protoplasts. Bright field and merged images are shown. (Scale bar = 5 μm).

To assess the subcellular localization of OsCAX2, we constructed vector expressing GFP-tagged rice OsCAX2 protein (*35S::CAX2-GFP*) and transiently expressed them in both of *Nicotiana benthamiana* leaves and rice protoplasts. The OsCAX2 protein was predominantly localized on the vacuoles in tobacco epidermal cells and in the rice protoplasts (Figure 2c and Supplementary Figure S1). The green fluorescence signal of OsCAX2-GFP overlapped neatly with the red fluorescence signal generated by the tonoplast-localized marker AtTPK1 in the rice protoplasts (Figure 2c). These results indicated that OsCAX2 is a vacuolar-localized protein.

### 3.2 *OsCAX2* mediates Cd sequestration into vacuole in yeast

To examine the Cd transport activity of OsCAX2, we expressed *OsCAX2* and its two truncated versions (^Δ^*NOsCAX2-16AA* and ^Δ^*NOsCAX2-40AA*) in yeast, since the phenomena of autoinhibition at the N-terminus of the *CAX* genes has been previously reported in other species (Kamiya et al., 2005). The *AtNRAMP1* gene from Col-0, a functional Cd transporter, was used as a positive control. On the SD-Ura medium without Cd, the yeast cells carrying the empty control, *AtNRAMP1, OsCAX2* and its truncated versions showed similar growth (Figure 3a). However, on SD-Ura medium containing Cd, the yeast cells expressing *AtNRAMP1* showed growth inhibition compared to the yeast cells expressing the empty control, while the yeast cells expressing *OsCAX2* and its truncated versions were similar to the control in growth (Figure 3a). The fluorescent signal of OsCAX2-GFP was partially localized with the signal of the tonoplast (Figure 3b). After 24 h Cd exposure, the *AtNRAMP1* and *OsCAX2* expressing cells had enhanced Cd accumulation compared to the empty control, and the Cd accumulation of the *OsCAX2*-expressing cells was 1.2-fold higher than that of the *AtNRAMP1*-expressing ones (Figure 3c), indicating that OsCAX2 has Cd transport activity without N-terminal autoinhibition. When loaded with Cd fluorescent probe, Leadmium™ Green AM, Cd treatment significantly enhanced the fluorescence intensity of OsCAX2-expressing protoplasts (Figure 3d and e). The fluorescence signal of OsCAX2-carrying cells was mainly localized in the vacuoles (Figure 3d), consistent with the vacuolar localization of OsCAX2 (Figure 3b). Taken together, these results demonstrated that OsCAX2 is a Cd transporter and may function in vacuolar intracellular sequestration of Cd in yeast.

**FIGURE 3.**
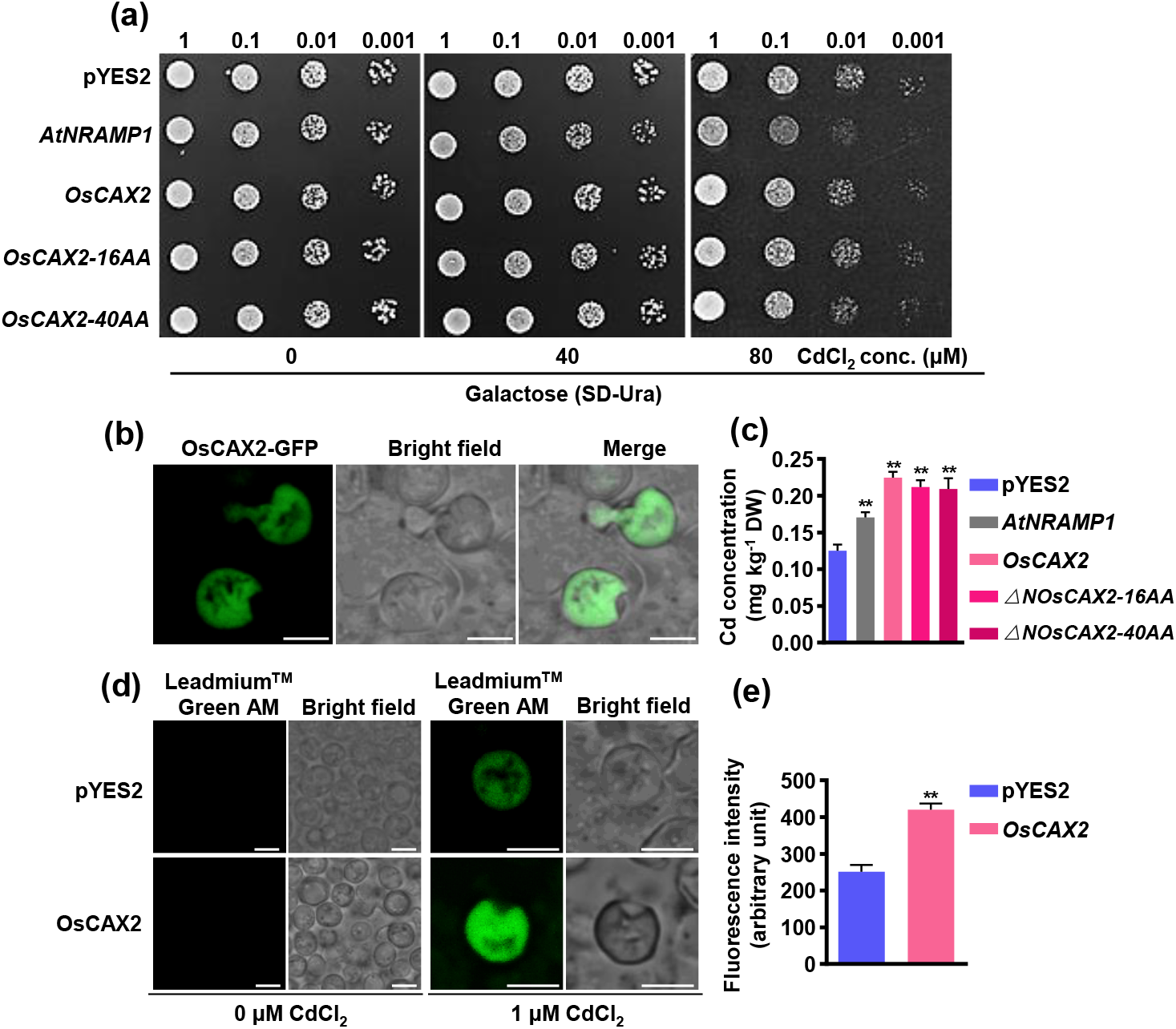
Cd transport activity and subcellular localization of *O*sCAX2 in yeast. (a) Growth of yeast strain BY4741 expressing *AtNRMAP1* (positive control), *OsCAX2, OsCAX2* truncated by 14 (Δ*NOsCAX2-16AA*) and 40 amino acids (Δ*NOsCAX2-40AA*) at the N-terminus or empty vector (pYES2) in SD-Ura medium containing different CdCl_2_ concentrations for 2 days at 30 °C. (b) Subcellular localization of OsCAX2 in yeast cells. Green color indicate the signals of OsCAX2-GFP (Scale bar = 5 μm). (c) Cd accumulation in yeast cells (BY4741) expressing *AtNRAMP1*, different versions of *OsCAX2* or empty vector after incubation in liquid medium containing 10 μM CdCl_2_ for 24 h. (de) Cd localization and quantification in the protoplasts of BY4741 yeast cells expressing *OsCAX2* or empty vector. Protoplasts were loaded with Leadmium™ Green AM for 60 min at 4 °C, and then incubated in solution with or without 1 μM CdCl_2_ for 5 min. (Scale bar = 5 μm). Fluorescence intensity was quantified for six protoplasts per sample using the Image J software (NIH, United States). Values are the mean ± SD of three independent replicates and statistical comparison was performed by ANOVA followed by planned comparison (\**P* < 0.05, \*\**P* < 0.01).

CAX transporters in most plant species have calcium (Ca) exchange activity and are able to complement the Ca sensitive phenotype of the yeast K667 mutant (Hirschi, 1999; Pittman and Hirschi, 2016; Shigaki and Hirschi, 2006). To test whether *OsCAX2* has the ability to suppress the Ca sensitivity of K667, the complete ORF of *OsCAX2* and its truncated version were cloned into K667 mutant along with empty vector as a control. The growth of the K667 cells expressing empty control, *OsCAX2* and truncated *OsCAX2* was all normal and very similar in the absence of Ca, but was severely inhibited in presence of 50 mM or 150 mM Ca (Supplementary Figure S2a). However, no differences were found between the cells of different transformants and the control was found under both of 50 mM and 150 mM Ca treatments (Supplementary Figure S2a), indicating that *OsCAX2* fails to complement the Ca sensitivity of K667. In the liquid medium containing 100 mM Ca, there were no differences in growth between the control and *OsCAX2*-carrying yeast cells (Supplementary Figure S2b), and the *OsCAX2*-expressing yeast cells had similar Ca accumulation to the control after 24 h Ca exposure (Supplementary Figure S2c). These findings demonstrated that *OsCAX2* has no Ca transport activity.

### 3.3 *OsCAX2* involves in Cd tolerance and root-to-shoot translocation at seedling stage in rice

To exam the role of *OsCAX2* in Cd tolerance and translocation we generated independent KO transgenic lines of *OsCAX2* using the CRISPR/Cas9 technology. The three KO lines selected for our study carried a 7-base deletion or a 1-base insertion, which led to a frame shift and the premature termination of *OsCAX2*, respectively (Figure 4a). We also generated OE lines by ectopically overexpressing *OsCAX2* driven by the *ubiquitin* promoter. The transcription levels of *OsCAX2* in the three OE lines chosen for our study were more than 22-fold higher than that in the WT (Figure 4b). The OsCAX2 protein levels of the WT, KO and OE lines were verified by western blotting (Figure 4c). To test the effect of *OsCAX2* on Cd tolerance, we compared the root and shoot growth of the WT, VC, KO and OE lines at seedling stage using hydroponic culture with five different Cd concentrations (0, 0.1, 1, 2 or 5 μM Cd) for 7 days. Since there was no significant difference between the WT and VC, only the WT was used as a control for comparative analysis. In the absence of Cd, the only significant difference was for the root length (RL) between the KO and WT lines with KO lines having longer RL (Figure 4d and e). In the presence of a low Cd concentration (0.1 or 1 μM), the only significant difference was for shoot dry weight (SDW) between the KO and WT lines under 1 μM Cd treatment with the KO lines having lower SDW (Supplementary Figure S3a and e). In the presence of 2 μM Cd, more significant differences were found between the transgenic lines and WT. The KO and OE lines had lower and higher SDW than the WT, respectively, the KO lines had shorter shoot length (SL) than the WT (Supplementary Figure S3a, c and e), and the OE lines had higher root dry weight (RDW) than the WT (Supplementary Figure S3d). At a higher Cd concentration (5 μM), all lines had reduced RL, SL, RDW, SDW and total dry weight (TDW) compared to the control (no Cd stress) with the KO and OE lines showing more and less reduction than the WT, respectively (Figure 4e-i). These results indicated that *OsCAX2* enhances Cd tolerance in rice.

**FIGURE 4.**
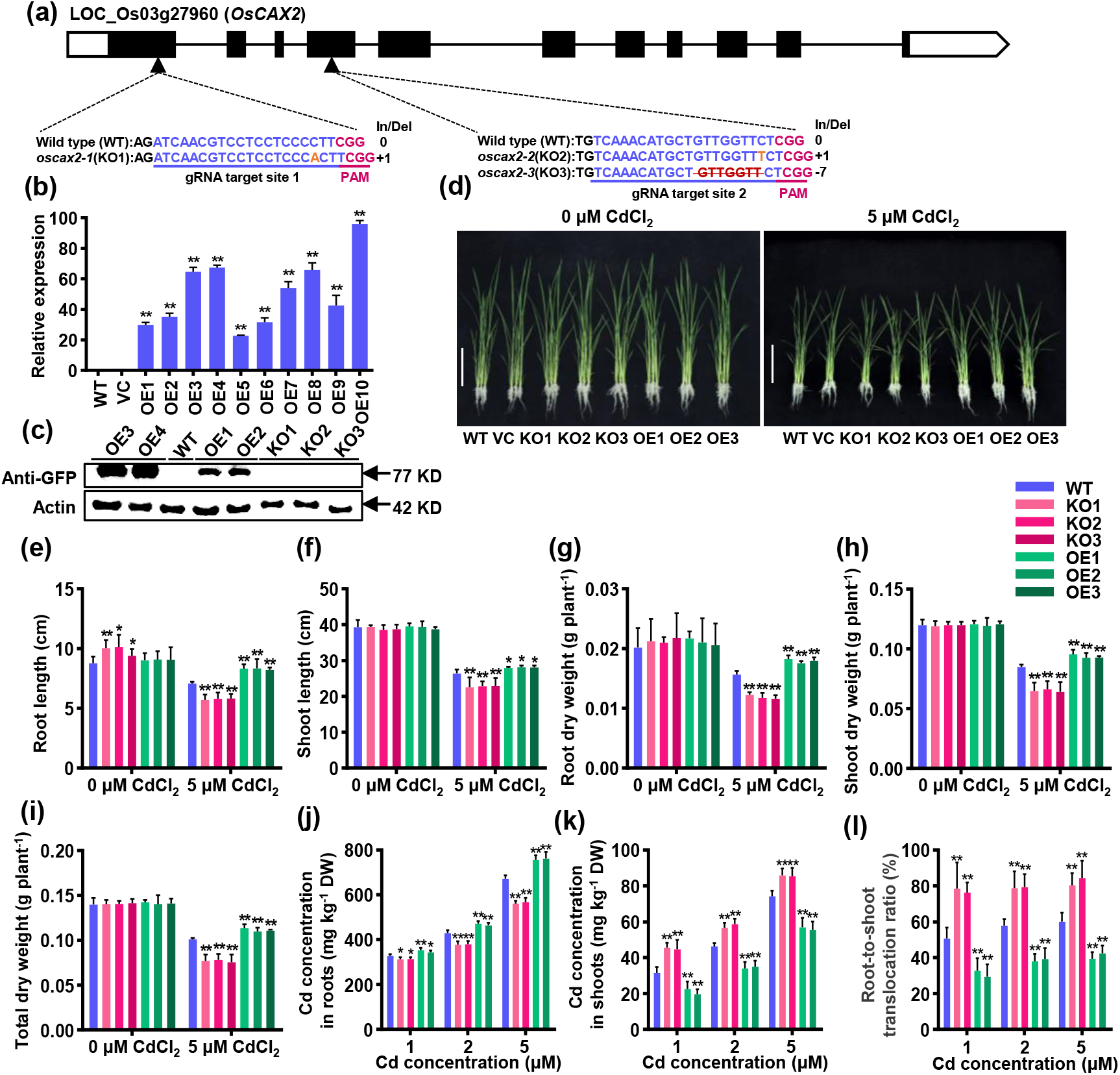
Phenotypic analysis of *OsCAX2* knockout (KO) mutant and overexpression (OE) lines. (a) Schematic representation of the gene model of *OsCAX2*. The arrow indicates the gRNA target site. The DNA sequences of the CRISPR/Cas9-edited *OsCAX2* gene. The 20-bp guide RNA (gRNA) spacer sequence for the Cas9/gRNA complex is in blue, and the protospacer adjacent motif (PAM) site is in red. Deleted nucleotides are depicted as dashes, and inserted nucleotides are shown in orange. The lengths of the insertions and/or deletions (In/Del) are shown on right. (b) Confirmation of *OsCAX2* overexpression in transgenic plants by RT-qPCR. The rice actin 1 gene (*Act1*) was used as an internal control. Data are means ± SD of three biological replicates. (c) Immunoblotting showing the proteins levels in the WT, KO (*oscax2-1, oscax2-2* and *oscax2-3*) and OE (OE1, OE2, OE3 and OE4) transgenic plants. (d) Phenotype of the WT, vector control (VC, segregated non-transgenic line), KO and OE lines (two-week-old) grown in hydroponic solution with or without 5 μM CdCl_2_ for 7 days. (Scale bar = 10 cm). (e-i) Quantitative phenotypic analysis. Root length (e), shoot length (f), root dry weight (g), shoot dry weight (h) and total dry weight (i). (j-k) Cd concentrations in roots (j) and shoots (k). (l) Cd root-to-shoot translocation ratio calculated as (shoot Cd content)/(root Cd content) × 100. Values are the mean ± SD of three independent replicates. The number of plants per line in a replicate is at least 6. Statistical comparison was performed by ANOVA followed by planned comparison (\**P* <0.05, \*\**P* <0.01).

To test whether *OsCAX2* is involved in Cd accumulation and translocation from roots to shoots, we analyzed the Cd accumulation in roots and shoots under different Cd concentrations (1, 2 or 5 μM CdCl_2_). Under these treatments, the root Cd concentration of the KO lines was significantly lower than that of the WT, whereas the root Cd concentration of the OE lines was significantly higher than that of the WT (Figure 4j). Contrastingly, the shoot Cd concentration of the KO and OE lines were significantly higher and lower than those of WT, respectively (Figure 4k). Compared with the WT, the KO lines had higher root-to-shoot Cd translocation while the OE had lower Cd translocation (Figure 4l). In the presence of 5 μM Cd, the KO lines had 15.5–16.6% lower root Cd concentration (Figure 4j) and 16.5–17.0% higher shoot Cd concentration than the WT (Figure 4k). In contrast, the OE lines had 12.4–13.4% higher root Cd concentration (Figure 4j) and 22.3–24.5% lower shoot Cd concentration than the WT (Figure 4k). Moreover, the KO lines had 33.6–42.9% higher Cd root-to-shoot translocation ratio (Figure 4l), while the OE lines had 29.5–37.0% lower translocation ratio than the WT (Figure 4l). We also measured the root and shoot concentrations of manganese (Mn), zinc (Zn), copper (Cu) and iron (Fe) under 5 μM Cd treatment, and found that they were similar in all lines (Supplementary Figure S4), indicating *OsCAX2* may have a relatively high Cd specificity. These results suggested that *OsCAX2* affects Cd accumulation and transport with its overexpression decreasing while its knockout increasing Cd concentration in shoots and root-to-shoot translocation.

### 3.4 *OsCAX2* facilitates Cd influx into rice roots at seedling stage

To evaluate the contribution of *OsCAX2* to Cd uptake in rice, we monitored Cd localization and measured Cd accumulation in roots using Leadmium™ Green AM. As expected, no Cd fluorescence signal was observed in all the tested lines under control (Figure 5a and b), whereas in the presence of Cd, the OE# and KO lines had obviously stronger and weaker signals than the WT, respectively (Figure 5a and b). Cd was highly accumulated in the meristematic zone and stele in the OE# lines (Figure 5a), consistent with the localization of OsCAX2 (Figure 2b. Furthermore, we conducted Cd uptake kinetics analysis. In the short-term (20 min) uptake experiment, Cd uptake of the OE lines was higher than that of the WT, which in turn was higher than that of the KO lines across the Cd concentration range from 0 to 5.0 μM (Figure 5c). Since apoplastic absorption is important in Cd uptake at 4 °C (Yan et al., 2016), net Cd uptake was calculated by subtracting the uptake at 4 °C from that at 25 °C to estimate the net symplastic uptake. The net uptake data fitted to the Michaelis–Menten kinetics equation very well with *R*^2^ being 0.983, 0.982 and 0.988 for the WT, KO and OE lines, respectively (Figure 5d). There were significant differences in the *Km* and *Vmax* values between the WT, KO and OE lines, with the OE lines having 66.6% higher *Km* and 52.4% higher *Vmax* values while the KO lines having 11.1% lower *Km* and 20.4% lower *Vmax* values than the WT (Figure 5d). We also measured the root Cd concentration of the WT, KO and OE lines subjected to 0 to 72 h treatment of 5.0 μM Cd. The data also fitted the Michaelis–Menten kinetic equation well with *R* being 0.957, 0.936 and 0.978, respectively (Figure 5e). Compared with WT, the OE lines had 252.1% higher *Km* and 162.3% higher *Vmax* values, while the KO lines had 42.5% lower *Km* and 41.7% lower *Vmax* values (Figure 5e). Taken together, these findings demonstrated that *OsCAX2* contributes to Cd influx into rice roots.

**FIGURE 5.**
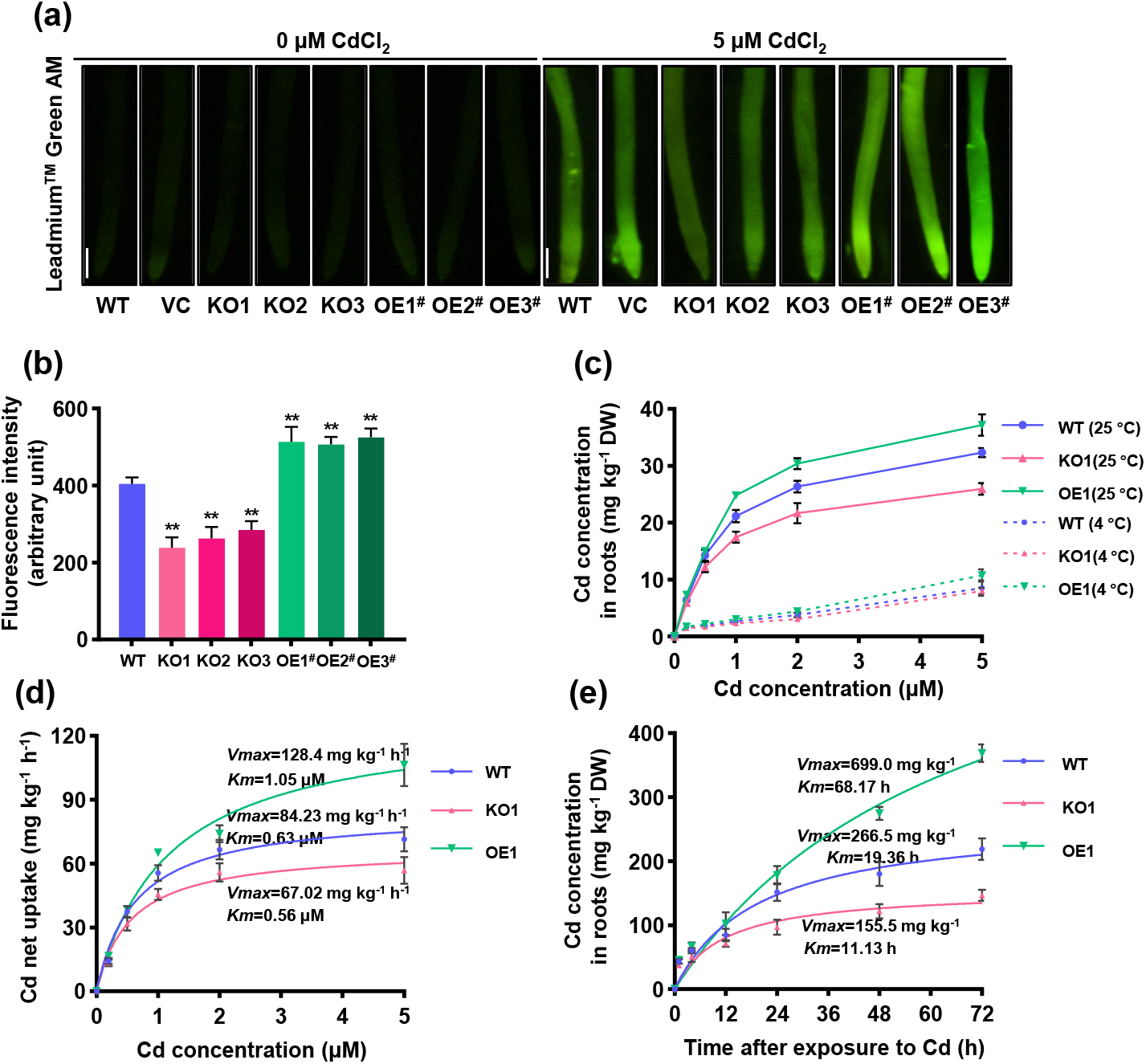
Effect of *OsCAX2* on Cd accumulation in roots at seedling stage. Three-week-old seedlings of the WT, KO and OE lines were used. (a) Visualization of Cd in the roots using Leadmium™ Green AM. Seedlings of the WT, KO and OE# (not labeled with GFP tag) lines were exposed to solution with or without 5 μM CdCl_2_ treatment for 7 days. (Scale bar = 500 μm). (b) The intensity of fluorescence. Six plants per line were measured using Image J software (NIH, United States). (c) Cd uptake at 4 °C and 25 °C. Seedlings were exposed to treatments of different CdCl_2_ concentrations at 4 °C or 25 °C for 20 min. (d) Cd net uptake calculated by subtracting the apparent uptake at 4 °C from that at 25 °C. (e) Time-course of Cd uptake. Seedlings were subjected to 5 μM CdCl_2_ treatment for indicated durations. Curves represent the fitted Michaelis-Menten equations, with *Km* and *Vmax* shown next to the curve. Values are the mean ± SD of three independent replicates. The number of plants per line in a replicate is at least 6. Statistical analysis was performed by ANOVA followed by planned comparison (\**P* < 0.05, \*\**P* < 0.01).

### 3.5 Overexpression of *OsCAX2* promotes Cd translocation from cytosol to vacuole and reduces Cd accumulation in xylem sap

To study the mechanism of *OsCAX2* in affecting Cd uptake and translocation we measured the Cd concentrations in the vacuoles and xylem sap of shoots. Firstly, we isolated protoplasts of the WT, KO and OE# lines, loaded them with Leadmium™ Green AM, and incubated them in the absence or presence of 1 μM CdCl_2_. When Cd was not added, the green fluorescence signal was not observed all lines (Figure 6a). The addition of 1 μM Cd increased the fluorescence signals in the protoplasts of all lines (Figure 6a). For the WT, the fluorescence signal was observed in the entire protoplast, indicating that Cd enters the cytoplasm and reaches the vacuole (Figure 6a). For the KO lines, the fluorescence signal was mainly in the cytoplasm with minimal being in the vacuole (Figure 6a). For the OE# lines, the fluorescence signal was mostly in the vacuole with minimal being in the cytoplasm (Figure 6a). To further verify the role of *OsCAX2* in sequestering Cd into vacuoles, we isolated the vacuoles from the protoplasts and measured the Cd accumulation in the vacuoles. The OE lines had significantly higher Cd accumulation in vacuoles than the WT, whereas the KO lines had significant lower Cd accumulation in vacuoles than the WT (Figure 6b). Taken together, these results suggested that *OsCAX2* promotes Cd sequestration into vacuoles.

**FIGURE 6.**
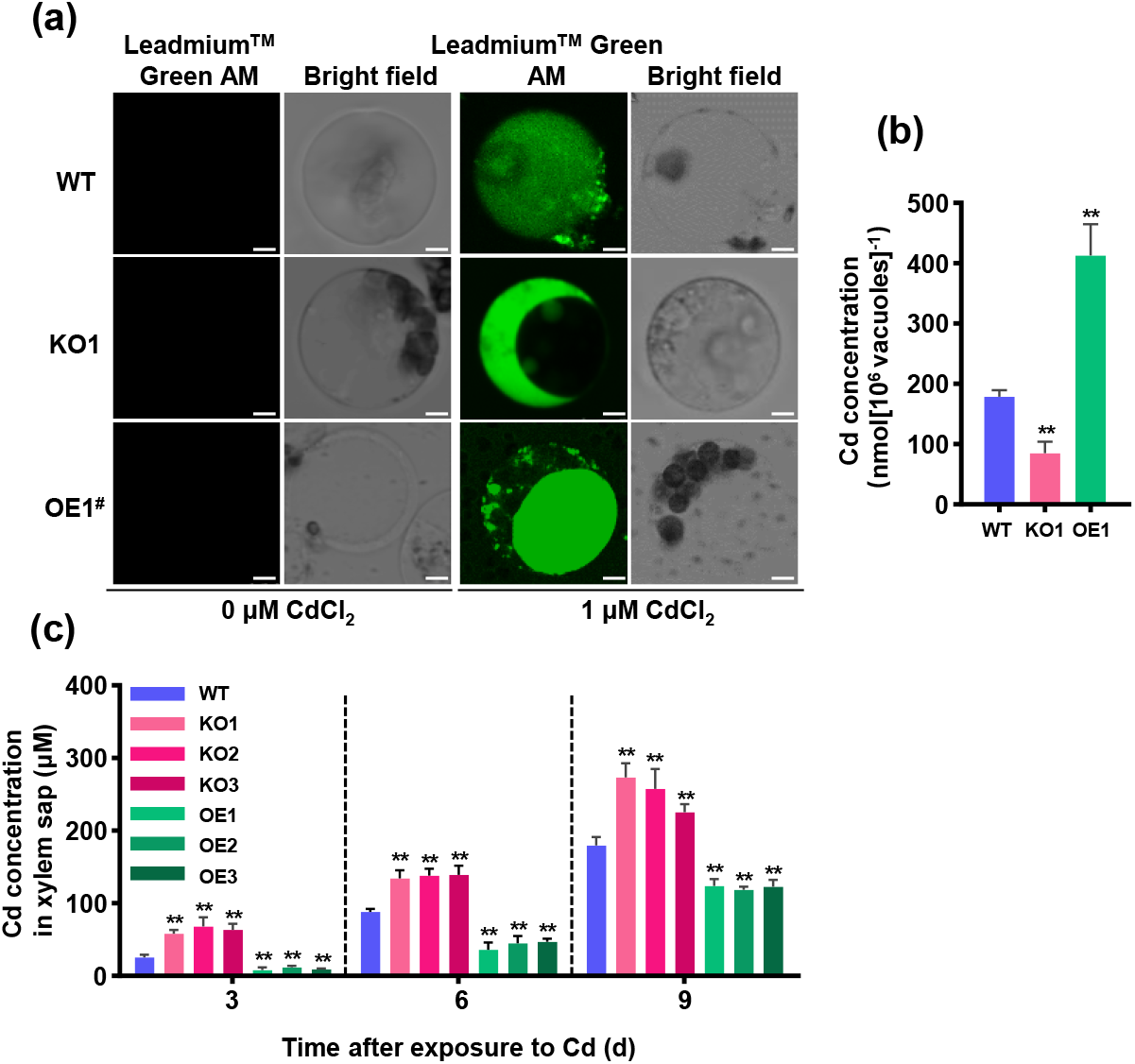
Cd concentrations in vacuoles and xylem sap. (a) Visualization of Cd accumulation in protoplasts. Protoplasts were loaded with Leadmium™ Green AM for 60 min at 4 °C and then incubated for 5 min in solution with or without 1 μM CdCl_2_. (Scale bar = 5 μm). (b) Cd concentration in vacuoles. Root vacuoles were extracted from the intact protoplasts of plants exposed to 5 μM CdCl_2_ for 7 days, and Cd concentration were determined by ICP-MS. (c) Cd concentration in xylem sap. Seedlings (three-week-old) were exposed to 5 μM CdCl_2_for 3, 6 or 9 days. Values are the mean ± SD of three independent replicates. The number of plants per line in a replicate is at least 6. Statistical analysis was performed by ANOVA followed by planned comparison (\**P* < 0.05, \*\**P* < 0.01).

We further measured the xylem sap Cd concentration in the shoots of the WT, KO and OE lines subjected to different durations (3, 6 or 9 days) of 5 μM Cd treatment. As expected, the xylem sap Cd concentration increased with treatment duration (Figure 6c). Significant differences between lines were found at all three points of measurements (Figure 6c), indicating that *OsCAX2* began functioning shortly after Cd treatment and continually functions during the exposure period. The largest differences between lines were found at the first measurement (3 days after Cd treatment), and at that time the KO lines had 125.4–165.4% higher while the OE lines had 53.2–69.3% lower xylem sap Cd concentration (Figure 6c).

Taken together, these results demonstrate that *OsCAX2* sequesters Cd into vacuole in roots for detoxification and reduces root-to-shoot translocation.

### 3.6 Overexpression of *OsCAX2* greatly reduces Cd accumulation in rice grains without negative effects on grain yield and agronomic traits

To test the effects of *OsCAX2* on Cd accumulation in rice grains, grain yield and important agronomic traits, we conducted a pot and a field experiment. The pot experiment was conducted with a soil Cd concentration of 1.5 mg kg^-1^ during the whole growth period. At harvest, plant height, effective tillers per plant, percentage of filled spikelet, 1000-grain weight, weight per panicle and yield per plant were not significantly different between the WT, KO and OE lines (Supplementary Figure S5a-f). The Cd concentration in brown rice of the WT was 0.22 mg kg^-1^, while that in the KO and OE lines was 0.43–0.46 and 0.010–0.018 mg kg^-1^, respectively (Supplementary Figure S5i). The Cd accumulation in brown rice of the OE lines was much lower than the maximum allowable limit (0.2 mg kg^-1^) set by FAO/WHO, while that in the KO and WT lines exceeded the limit (Supplementary Figure S5i). The KO lines had 53.3–69.7% and 47.1–67.3% higher Cd concentration in straws and leaves, whereas the OE lines had 83.1–85.5% and 73.5–81.7% lower Cd concentration in straws and leaves (Supplementary Figure S5g and h).

For the field experiment, the soil Cd concentration of was 1.8 mg kg^-1^ (pH at 5.5). Similar to the pot experiment, there were no differences between lines in growth and development (Figure 7a), grain yield and other agronomic traits at maturity (Figure 7b-g). Compared with WT, Cd accumulation in brown rice significantly increased by 77.4–81.0% in the KO lines and decreased by 70.0–78.1% in the OE lines, respectively (Figure 7h). The Cd concentration in brown rice of the OE lines was 0.053–0.062 mg kg^-1^, much lower than the FAO/WHO limit of Cd in rice grains (Figure 7h). These results suggested that overexpressing *OsCAX2* greatly reduces Cd accumulation in rice grains with no effects on grain yield and other important agronomic traits under different growing environments.

**FIGURE 7.**
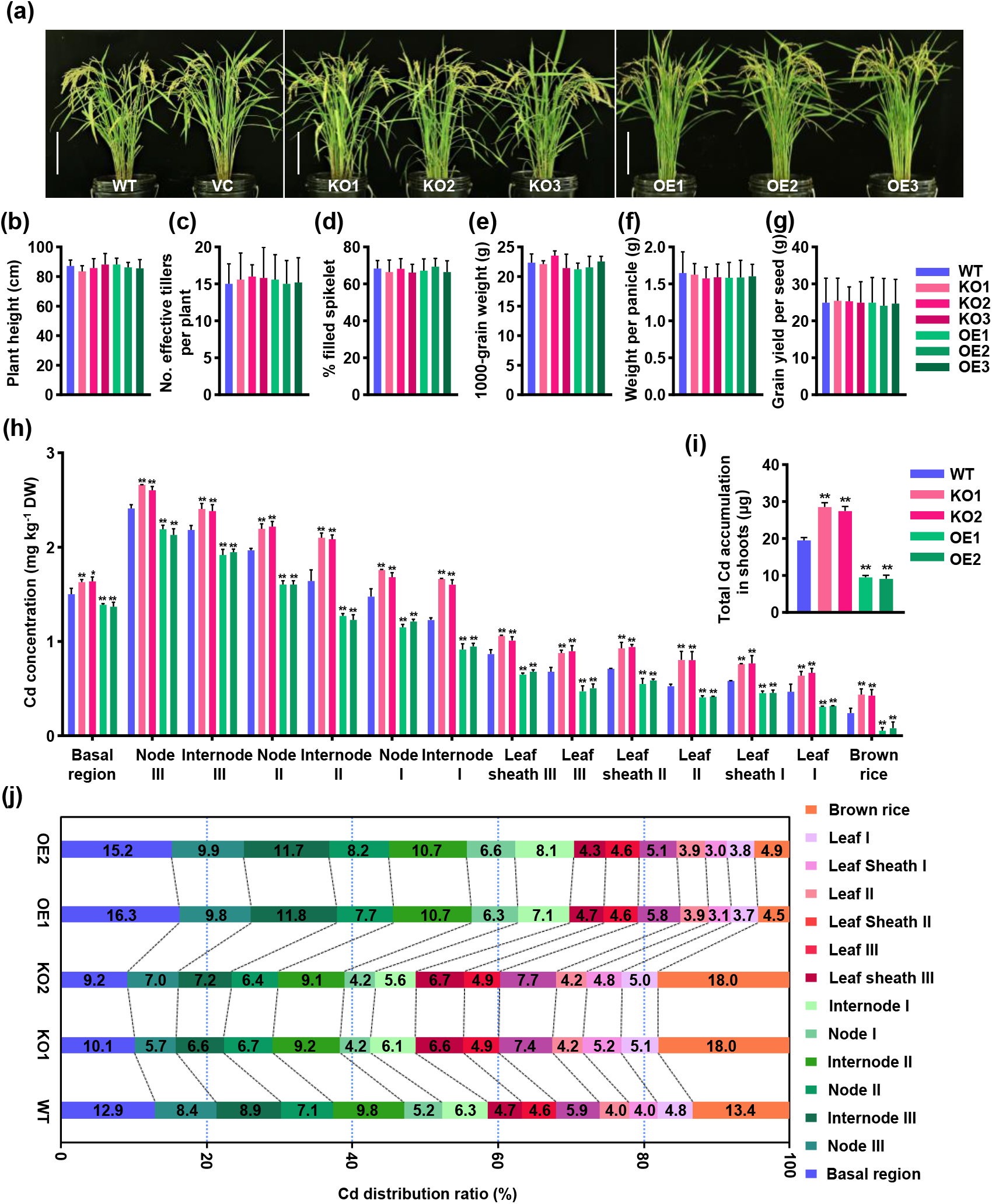
Effects of *OsCAX2* on Cd distributions in different tissues/organs and agronomic traits at mature stage. (a) Phenotype of the WT, KO and OE lines grown in a paddy field with Cd content 1.8 mg kg^-1^ and pH at 5.5. (Scale bar = 20 cm). (b-g) Agronomic traits. Plant height (b), Number of effective tillers per plant (c), % filled spikelet (d), 1000-grain weight (e), weight per panicle (f) and grain yield per seed (g). (h) Cd concentrations of different tissues/organs. (i) Total above-ground Cd content calculated as the sum of the contents in all above-ground tissues/organs. (j) Cd distribution ratio calculated as (tissue Cd content)/(total above-ground Cd content) × 100. Values represent means ± SD of biological replicates (n = 6). Statistical analysis was performed by ANOVA followed by planned comparison with WT(\**P* <0.05, \*\**P* <0.01).

### 3.7 *OsCAX2* mediates Cd accumulation and distribution in different aboveground tissues/organs at maturity under field conditions

To investigate the effects of *OsCAX2* on Cd accumulation and distribution in different aboveground tissues/organs at maturity, we analyzed Cd concentration in the basal regions, nodes I-III, internodes I-III, leaf sheaths I-III, leaves I-III and brown rice of plants grown in the field experiment. The Cd concentrations in all tissues were significantly higher in the KO lines but significantly lower in the OE lines than those in the WT (Figure 7h). Compared to WT, the total Cd accumulation in the KO lines was 41.0–46.4% higher, whereas that in the OE lines was 51.3–53.1% lower (Figure 7i), indicating that *OsCAX2* plays an important role in the total aboveground Cd accumulation. The basal regions, nodes I-III, internodes I-III, leaves I-III, leaf sheaths I-III and brown rice respectively accounted for 12.9%, 20.7%, 25.0%, 14.6%, 13.4% and 13.4% of the total aboveground Cd content in the WT (Figure 7j), whereas the corresponding proportions were 9.2–10.1%, 16.6–17.6%, 21.9%, 19.2%, 14.1–14.3% and 18.0% in the KO lines and were 15.2–16.3%, 23.8–24.7%, 29.6–30.5%, 12.4–13.6%, 12.2–12.3% and 4.5–4.9% in the OE lines (Figure 7j), suggesting that *OsCAX2* also greatly affects the Cd distribution to different aboveground tissues/organs. Taken together, these results demonstrated that *OsCAX2* plays important roles in both of the total aboveground Cd accumulation and the distribution to different tissues/organs, and the low grain Cd accumulation in the OE lines is caused by reduction in the total accumulation and distribution to grains.

To investigate the effects of *OsCAX2* on contents of other elements at mature stage, we measured Mn, Cu, Zn and Fe contents in straws, leaves and brown rice. There were no significant differences between all lines in the concentrations of Mn and Zn in the straws and leaves (Supplementary Figure S6a-d). However, the OE and KO lines had significantly higher and lower brown rice concentrations of Mn and Zn than the WT, respectively (Supplementary Figure S6e and f). The Cu concentration in the straws was similar in all lines (Supplementary Figure S6g), whereas the Cu concentrations in the leaves and brown rice were significantly lower in the OE lines and higher in the KO lines than those in the WT, respectively (Supplementary Figure S6h and i). Compared to the WT, the KO and OE lines had significantly lower and higher Fe concentrations in the straws, leaves and brown rice, respectively (Supplementary Figure S6j-l). Taken together, these results suggested that *OsCAX2* affects the homeostasis of other metal ions in rice, and the OE lines have reduced accumulation of Cd and Cu but enhanced accumulation of Mn, Zn and Fe in grains. These results and the results on grain yield and agronomic traits (Figure 7a-g and Supplementary Figure S6a-f) demonstrated that *OsCAX2* is an ideal gene for developing Cd safety rice varieties via transgenic approach.

## 4 DISCUSSION

To facilitate the development of Cd-safe rice varieties, extensive studies using reverse and forward genetic approaches have been conducted to identify genes involving in Cd tolerance and accumulation in rice and other crops. Although a few genes including *OsNRAMP5* and *OsHMA3* have been shown to be very promising (Yan et al., 2016; Yang et al., 2021), more genes with large effect on grain Cd content without or with little adverse effect on other traits are needed. In this study, by systematically characterizing the function of *OsCAX2* in Cd tolerance and accumulation in rice and investigating the possible mechanisms underlying the function, we reported that OsCAX2 is an important novel Cd transporter ideal for developing Cd-safe varieties via transgenic approach.

OsCAX2 is located on vacuoles and has Cd transport activity in yeast (Figure 3). In rice, OsCAX2 is vacuole-localized (Figure 2c) and very responsive to Cd stress and expresses more strongly in roots than in shoots (Figure 1) with strongest expression being seen in the root exodermis, parenchyma in cortex, endodermis and stele cells (Figure 2b). *AtCAX1* is highly expressed in leaf and floral tissues (Cheng et al., 2005). *AtCAX2* is strongly expressed in the leaf and vascular tissues and in root tip (Pittman et al., 2004). *AtCAX3* is mainly expressed in root, especially in root tip (Cheng et al., 2005). *AtCAX4* is mainly expressed in the apical and lateral root primordial (Mei et al., 2009). The expression of *AtCAX4* is induced by Cd stress (Mei et al., 2009). Exposing barley (*Hordeum vulgare*) plants to Cd stress changes the expression of *CAX* genes (Schneider et al., 2009).

We found that *OsCAX2* greatly contributes to Cd tolerance and accumulation (Figure 4d-l). Overexpressing *OsCAX2* enhances root Cd uptake and content (Figure 4j and 5), and reduces shoot Cd content (Figure 4 k), and Cd root-to-shoot translocation (Figure 4l). *OsCAX2* contributes significantly to Cd transport in roots by reducing xylem sap Cd concentration through sequestering Cd to vacuoles (Figure 6). Overexpressing *OsCAX2* reduces the brown rice Cd content by more than 70% due to the great reduction in the total aboveground Cd accumulation and distribution to grains (Figure 7h and Supplementary Figure S5i). In *Arabidopsis*, four *CAX* genes have been showed to involve in Cd tolerance and transport with AtCAX2 and AtCAX4 showing very strong Cd transport capabilities due to a higher Cd selectivity (Koren’kov et al., 2007). Expression of *AtCAX2* or *AtCAX4* in tobacco roots reduces Cd translocation to the aboveground parts by enhancing sequestration of the metal in the roots (Korenkov et al., 2009). *SaCAX2h* encoding a CAX2-like transporter isolated from *Sedum alfreddi*, a Zn/Cd hyperaccumulating plant, enhances Cd accumulation in tobacco (Zhang et al., 2016). *AtCAX1* overexpressed in petunia (*Petunia hybrida*) plants confers increased tolerance to Cd (Wu et al., 2011). Loss-of-function of *AhCAX1* results in a high sensitivity to Cd but only at low concentration of Ca (Baliardini et al., 2015). However, the role of *AtCAX1* in Cd tolerance is likely via maintaining cytosolic Ca levels to avoid ROS accumulation instead of effecting on Cd transport (Baliardini et al., 2015). The transgenic tobacco and potato plants overexpressing *AtCAX1* show Ca deficiency disorders such as yellow leaves and dwarf plants (Hirschi, 1999; Zorrilla et al., 2019). The function of *OsCAX2* in Cd tolerance and transport in our study was similar to that reported for *AtCAX2*, which is not unexpected since OsCAX2 is tonoplast-localized and highly homologous to AtCAX2 (Kamiya et al., 2005; Zou et al., 2021). *OsCAX2-*mediated tolerance does not seem to be Ca-dependent (Figure 4d and Supplementary Figure S2). Similarly, *AtCAX2* has no response to Ca (Zheng et al., 2021). Although all *Arabidopsis* CAX transporters are tonoplast-localized, OsCAX3 and OsCAX4 in rice are plasma member-located (Qi et al., 2005; Zou et al., 2021). The rice CAX transporters may transport Cd using the tonoplast-localized ones to remove Cd from the cytosol and releases H^+^, and the plasma membrane-located ones to efflux H^+^ from the extracellular space and releases Cd. Clearly, more studies are needed to investigate the roles of other rice *CAX* genes including how they work together to achieve Cd detoxification in rice.

The vacuole is a pivotal organelle functioning in temporary storage for essential and toxic metabolites, mineral nutrients and toxic pollutants in plants (Peng and Gong, 2014). In the present study, we showed that the major mechanism for *OsCAX2*-mediated Cd tolerance and accumulation is the sequestration Cd into vacuoles, which reduces Cd root-to-shoot translocation (Figure 4l and 6). Only a few tonoplast-located Cd transporters, including OsHMA3, OsNRAMP2, OsABCC9 and OsMTP1 were functionally validated in rice previously. *OsHMA3, OsABCC9* and *OsMTP1* mediate the transport of Cd from the cytoplasm into the vacuole and thus the root-to-shoot translocation of Cd (Menguer et al., 2013; Sasaki et al., 2014). *OsNRAMP2* mediates Fe remobilization and affects translocation of Cd from vegetative tissues to rice grains (Chang et al., 2022; Zhao et al., 2018). Among them, *OsHMA3* had the largest effect in Cd tolerance and translocation. Overexpression of *OsHMA3* enhances root Cd accumulation, decreases shoot Cd accumulation and root-to-shoot translocation, and dramatically reduces grain Cd content in rice (Sasaki et al., 2014). The natural variations in the coding regions of *OsHMA3* contributes to the phenotypic difference between rice varieties with the dysfunction alleles conferring increasing grain Cd content (Miyadate et al., 2011; Ueno et al., 2011; Ueno et al., 2010; Yan et al., 2016). Natural variations in the promoter of *OsHMA3* have recently been showed to contribute to the differences between varieties as well (Liu et al., 2020). Our results on *OsCAX2* OE lines are very similar to the reported results on *OsHMA3* OE lines. In addition, heterologous expression of *OsHMA3* and *OsMTP1* in tobacco demonstrated that they mediate Cd tolerance through the phytochelatin (PC) and ROS synthesis pathways, respectively (Cai et al., 2019; Das et al., 2016). Our unpublished results also showed that *OsCAX2* mediates Cd tolerance through both the ROS and PC pathways. However, *OsHMA3* mainly expresses in roots, while *OsCAX2* expresses in roots, leaves and nodes, although they have similar localization in the root cells. Furthermore, *OsHMA3* is not responsive to Cd, while *OsCAX2* is strongly upregulated by Cd. It is worthy to note that nonsynonymous natural mutations were not found for *OsCAX2* in more than 4000 germplasm with high quality sequences (Data not shown). Further work may prove that the combined use of these two transporters is a good strategy for developing Cd-safety rice varieties. Non-vacuole-localized transporters also play roles in Cd sequestration in rice. For instance, the trans-Golgi network localized OsMTP11 transports Cd and other heavy metals into the vacuoles (Zhang and Liu, 2017). The ER-located OsLCT2 reduces Cd accumulation in rice shoots and grains by limiting the xylem loading of Cd and restricting the root-to-shoot Cd translocation (Tang et al., 2021).

*OsCAX2* was found to promote root Cd uptake (Figure 5). This is undesirable for reducing grain Cd content, since it counterbalances its desirable effect on root-shoot translocation. Among the metal transporters involved in Cd uptake in rice, only *OsNRAMP5* appears to have a larger effect than *OsCAX2*. The effect of *OsNRAMP5* is so great that Cd-safe lines can be developed by knocking it out or down. However, the knockout of *OsNRAMP5* may also have negative effects on other metal elements, particularly Mn, and thus reduces grain yield (Sasaki et al., 2012; Yang et al., 2019). The *OsCAX2* OE lines had increased Mn accumulation and no adverse effect on grain yield and other important agronomic traits (Figure 7a-g and Supplementary Figure S6e). Therefore, simultaneously knocking out *OsNRAMP5* and overexpressing *OsCAX2* may be better strategy for achieving Cd safety and stable production. Additionally, whether and how *OsCAX2* and *OsNRAMP5* work together will be an interesting topic for future study.

We also found that the *OsCAX2* contributes to Cd distribution to different tissues/organs at maturity (Figure 7j), and *OsCAX2* reduces grain Cd accumulation partially through reducing Cd distribution to grains, for which the nodal transport may play a critical role. Among the reported node-expressed transporters accountable for Cd transport, the plasma membrane-located OsHMA2 is well-studied, and it acts as an efflux transporter of Cd, responsible for Cd loading to xylem (Satoh-Nagasawa et al., 2012). At the reproductive stage, *OsHMA2* expresses in nodes and is involved in reloading Cd from the intervening parenchyma tissues into the phloem of the diffuse vascular bundles (Yamaji et al., 2013). Unfortunately, the knockout of *OsHMA2* reduces grain yield due to the accompanied reduction in Zn (Yamaji et al., 2013). The plasma membrane-located OsLCT1 functions as a Cd efflux transporter, and involves in loading Cd into phloem sieve tube (Uraguchi et al., 2011). *OsLCT1* expresses around the enlarged vascular bundles and diffuse vascular bundles in the node at the reproductive stage and knockdown of *OsLCT1* decreases Cd concentration in rice grains without affecting mineral nutrient concentrations and plant growth (Uraguchi et al., 2011). *OsCCX2* is expressed in xylem, functions in loading Cd into xylem vessel, and mediates direct root-derived Cd transport to grain (Hao et al., 2018). The knockout of *OsCCX2* significantly decreases both of grain Cd content and yield. *OsZIP7* involves in xylem-loading in roots and intervascular transfer in nodes to deliver Cd upward in rice (Tan et al., 2019). It is worth mentioning that *OsNRAMP2* involves in Cd distribution from leaves and straws to rice grains, although it was not expressed at nodes (Chang et al., 2022). Clearly, both of xylem- and phloem-mediated Cd transport activities are important in determining grain Cd level. Although systematic comparison between these genes has not been done yet, it seems that the effects of *OsHAM2* and *OsCCX2* are larger. Compared to *OsHAM2* and *OsCCX2*, *OsCAX2* has the advantage that it does not have adverse effects on grain yield and other agronomic traits.

## 5 CONCLUSIONS

In summary, through comprehensive experiments we demonstrated that the tonoplast-localized *OsCAX2* plays a key role in Cd tolerance, translocation and accumulation in rice. It functions in root uptake, vacuole sequestration, xylem loading, and tissue distribution of Cd. Importantly, overexpressing *OsCAX2* greatly reduces grain Cd accumulation in rice without adverse effects on yield and agronomic traits, which is ideal for the developing Cd-safe varieties via transgenic approach. Future studies in deciphering the molecular mechanisms of *OsCAX2-mediated* Cd tolerance and accumulation and the possible direct or indirect interactions of OsCAX2 with other Cd transporters such as OsNRAMP5, OsHMA3 and OsHMA2 in rice will provide more solid foundation for designing efficient breeding strategies.

## Supporting information

OsCAX2 Supplementary Figures

## ACKNOWLEDGEMENTS

This work was supported by the National Key R&D Program of China (2020YFE0202300), the Shenzhen Science and Technology Program (JCYJ20200109150650397 and JCYJ20190813104211014) and the Agricultural Science and Technology Innovation Program Cooperation and Innovation Mission (CAAS-XTCX2016001).

## AUTHOR CONTRIBUTIONS

G.Y. and J.C. conceived the idea and supervised this study. W.Z., J.Z., Y.C., D.C. and M.Z. performed the experiments. W.Z. analyzed the data. H.H., J.Z., L.M., J.C., D.C., Y.C and M.Z. participated in the result analysis. W.Z. and G.Y. wrote the manuscript.

## CONFLICT OF INTEREST

The authors declare no conflicts of interest.

## DATA AVAILABILITY

All data supporting the findings of this study are available within the paper and within its supplementary data published online.

## Notes

### Competing Interest Statement

The authors have declared no competing interest.

