## Supplementary material for "The Cation/H^+^ exchanger OsCAX2 is involved in cadmium tolerance and accumulation through vacuolar sequestration in rice": OsCAX2 Supplementary Figures

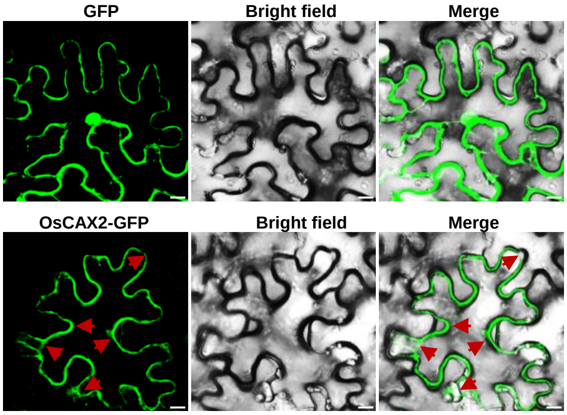


**FIGURE S1** Subcellular localization of OsCAX2 in *N. benthamiana* leaf cells. The red arrows point to the signal of tonoplast-localized. 35S::GFP, the positive control. (Scale bar = 10 μm).

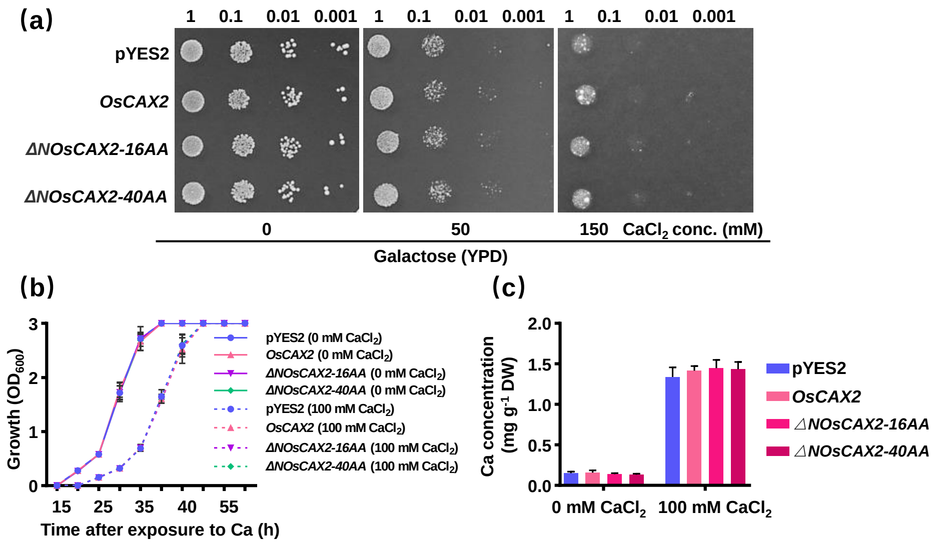


**FIGURE S2** The effect of *OsCAX2* on Ca tolerance and accumulation in yeast. (a) Growth of yeast strain K667 expressing empty vector (pYES2), *OsCAX2* and *OsCAX2* truncated versions in YPD medium containing different Ca concentrations (0, 50 or 100 mM). The plates were incubated at 30 °C for 2 days. (b) Growth of yeast cells carrying empty vector or *OsCAX2* in liquid medium with or without 100 mM $CaCl_2$. (c) Ca accumulation in yeast strain K667 expressing empty vector or O*sCAX2* in liquid medium with or without 100 mM $CaCl_2$ for 24 h. Values are the mean ± SD of three independent replicates and statistical analysis was performed by ANOVA followed by planned comparison with the K667 expressing empty vector.

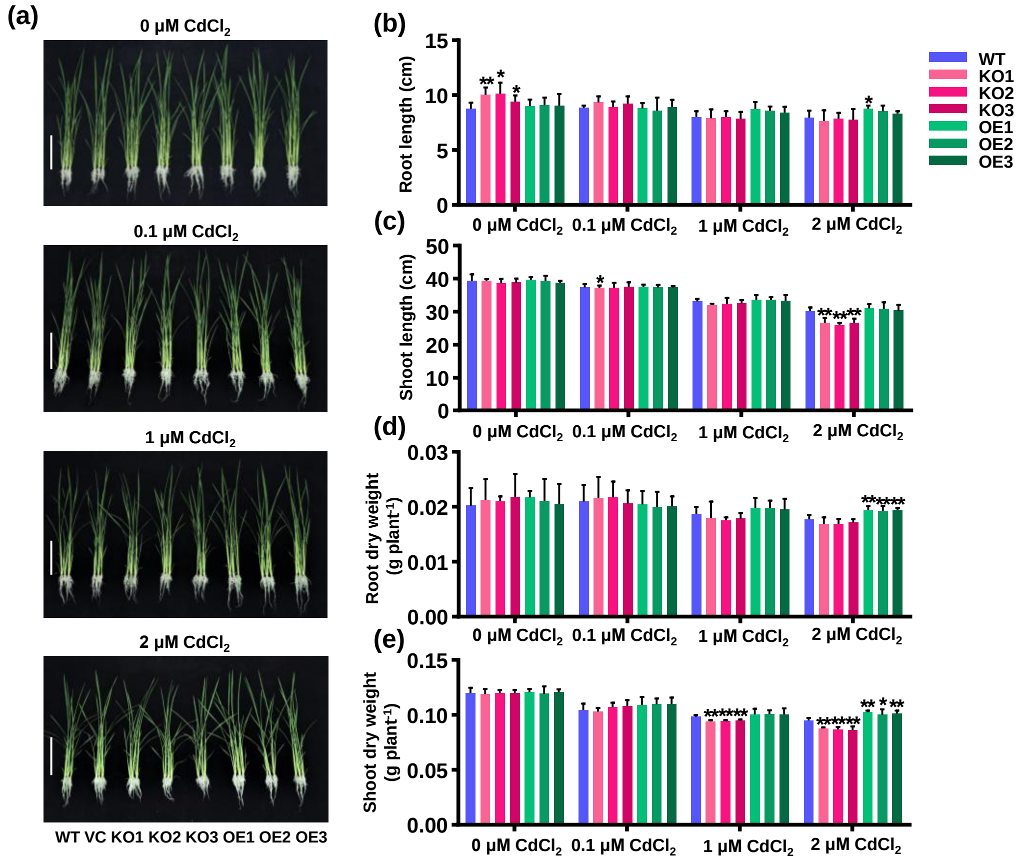


**FIGURE S3** Effect of *OsCAX2* on growth at seedling stage. (a) Phenotype of the WT, vector control (VC, segregated non-transgenic line), KO and OE lines (two-week-old) grown in hydroponic solution with 0, 0.1, 1 and 2 μM $CdCl_2$ for 7 days. (Scale bar = 10 cm). (b-e) Phenotypic analysis. Root length (b), shoot length (c), root dry weight (d), shoot dry weight (e) of all lines. The Cd-free (0 μM) treatment in this experiment was consistent with that in Figure 4. Values are the mean ± SD of three independent replicates each containing at least 6 plants per line and statistical analysis was performed by ANOVA followed by planned comparison (**P* < 0.05, ***P* < 0.01).

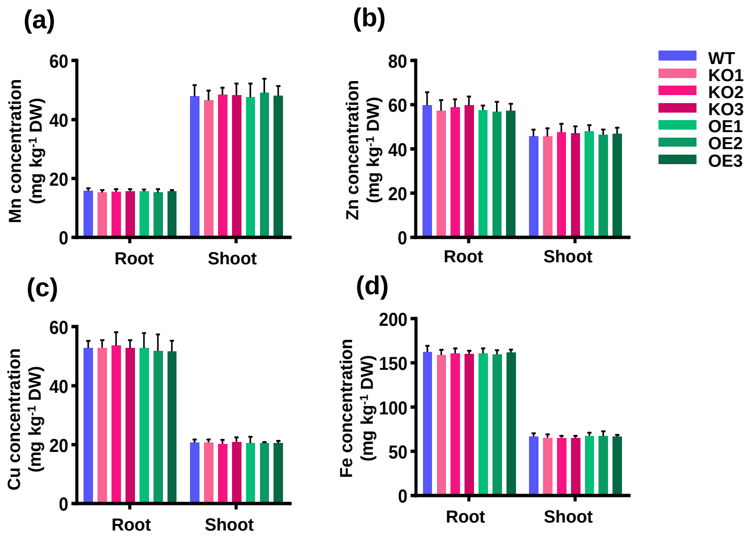


**FIGURE S4** Effect of *OsCAX2* on Mn, Zn, Cu and Fe concentrations at seedling stage. Seedlings of the WT, KO and OE lines (two-week-old) were exposed to 5 μM $CdCl_2$ for 7 days. (a-d) Mn, Zn, Cu and Fe concentrations in roots and shoots. Values are the mean ± SD of three independent replicates each containing at least 6 plants per line and statistical analysis was performed by ANOVA followed by planned comparison with WT.

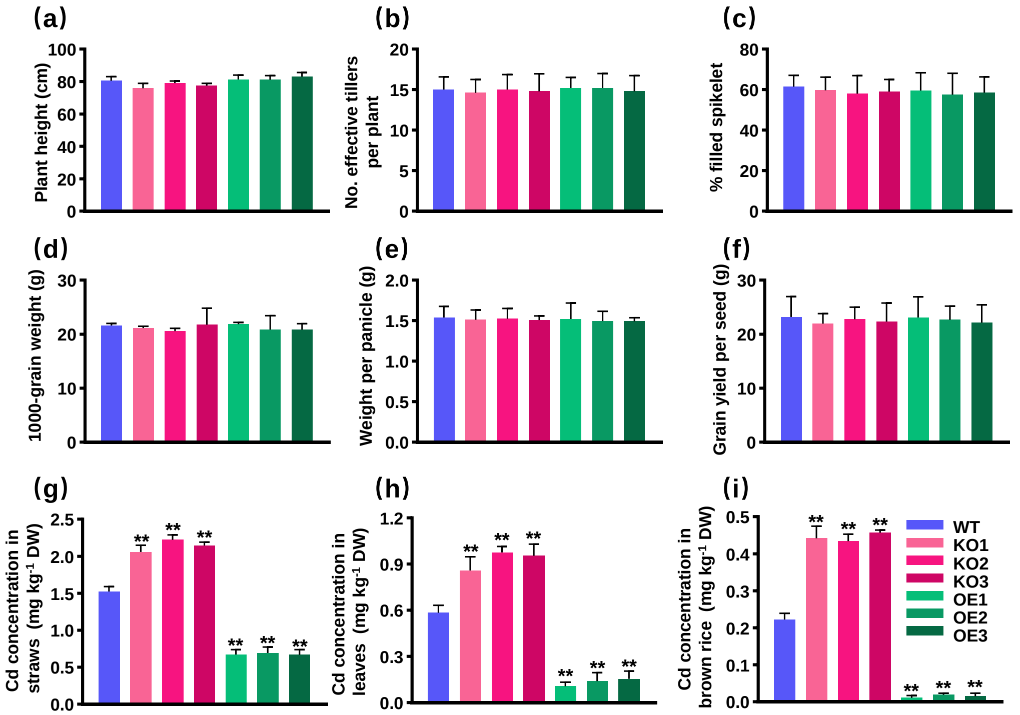


**FIGURE S5** Effects of *OsCAX2* on Cd accumulation and agronomic traits at maturity in a pot experiment. (a-f) Agronomic traits. Plant height (a), No. effective tillers per plant (b), % filled spikelet (c), 1000-grain weight (d), weight per panicle (e) and grain yield per seed (f). (g-i) Cd concentration. Cd concentrations in straws (g), leaves (h) and brown rice (i). Values represent means ± SD of biological replicates (n = 6). Statistical analysis was performed by ANOVA followed by planned comparison with WT (**P* < 0.05, ***P* < 0.01).

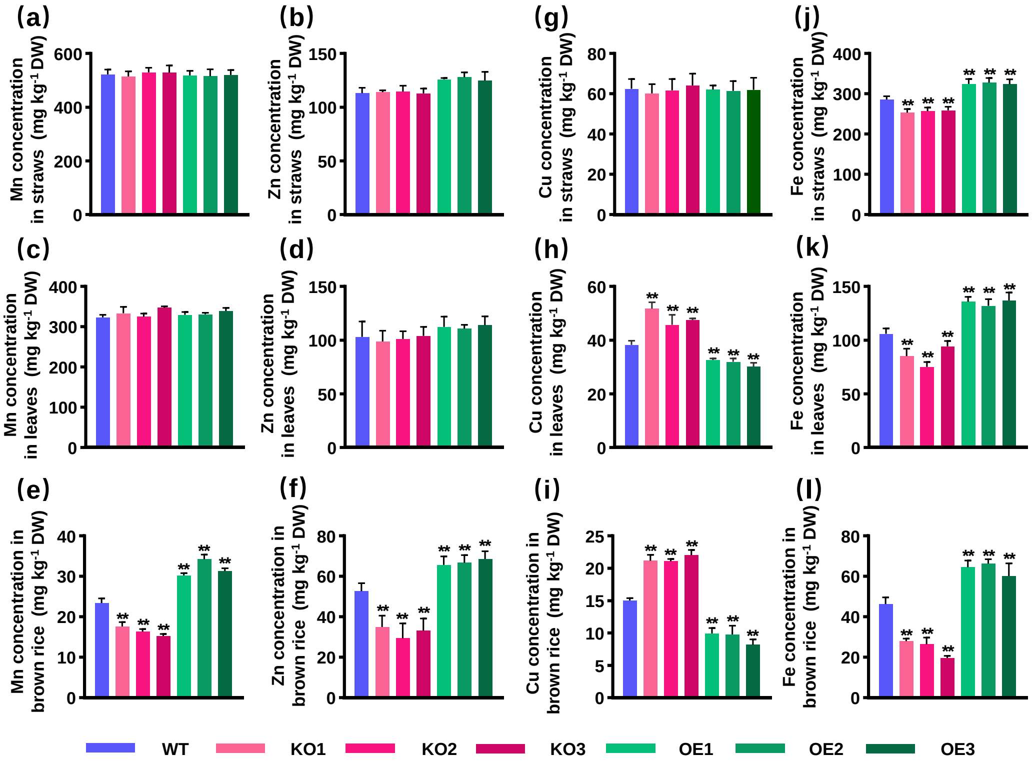


**FIGURE S6** Effect of *OsCAX2* on Mn, Zn, Cu and Fe concentrations at mature stage in a field experiment. A contaminated paddy field with soil Cd content 1.8 mg kg$^{-1}$ and pH at 5.5 in Shenzhen, China. (a-l) Mn, Zn, Cu and Fe concentrations in straws, leaves and brown rice. Values represent means ± SD of biological replicates (n = 6). Statistical analysis was performed by ANOVA followed by planned comparison with WT (*$P$ < 0.05, **$P$ < 0.01).
